# Failure to clear developmental apoptosis contributes to the pathology of RNASET2-deficient leukoencephalopathy

**DOI:** 10.1101/744144

**Authors:** Noémie Hamilton, Holly A. Rutherford, Hannah M. Isles, Jessica J. Petts, Thomas Weber, Marco Henneke, Jutta Gärtner, Mark Dunning, Stephen A. Renshaw

## Abstract

The contribution of microglia in neurological disorders is emerging as a leading driver rather than a consequence of pathology. RNAseT2-deficient leukoencephalopathy is a severe childhood white matter disorder affecting patients in their first year of life and mimics a cytomegalovirus brain infection. The early onset and resemblance of the symptoms to an immune response suggest an inflammatory and embryonic origin of the pathology. In this study, we identify deficient microglia as an early marker of pathology. Using the *ex utero* development and the optical transparency of an *rnaset2*-deficient zebrafish model, we found that dysfunctional microglia fail to clear apoptotic neurons during brain development. This was associated with increased number of apoptotic cells and behavioural defects lasting into adulthood. This zebrafish model recapitulates all aspect of the human disease to be used as a robust preclinical model. Using microglia-specific depletion and rescue experiments, we identified microglia as potential drivers of the pathology and highlight tissue-specific approaches as future therapeutic avenues.

## Introduction

Developmental apoptosis in the CNS allows the brain to eliminate excess neurons produced during development (Meier *et al.*, 2000; Nijhawan *et al.*, 2000; Dekkers *et al.*, 2013). Dying neurons are cleared by the main phagocyte in the CNS—the microglia—and this immune response is necessary to allow natural clearing of apoptotic neurons. However, disordered immunity in the brain is also a feature of many neurological disorders and plays a crucial role in the onset and progression of many neurodegenerative and monogenic white matter diseases.

White matter disorders such as leukodystrophies are characterised by myelin defects and white matter degeneration. The early onset of clinical symptoms includes loss of motor function and cognitive decline, with patients often dying at a young age (Bielschowsky and Henneberg, 1928). Patients as young as a few months old can present with neurological deficits, suggesting a foetal origin of the disease. At least 200 rare leukodystrophies have been identified to date, many of which are caused by the dysfunction of oligodendrocytes and subsequent loss of myelin integrity (United Leukodystrophy Foundation, 2019). More recently, the contribution of microglia and neuroinflammation to disease pathology has been recognised, prompting reclassification of these diseases and a requirement for further understanding of the mechanisms underlying disease progression (Kevelam *et al.*, 2016; van der Knaap and Bugiani, 2017).

Neuroinflammation is a major contributor to the progression of leukodystrophies, with increased immune activity detected even before the onset of myelin loss. Evidence of microglial activation preceded the reduction of myelin viability in post mortem brains of patients suffering from metachromatic leukodystrophy (MLD) and X-linked adrenoleukodystrophy (X-ALD) (Bergner *et al.*, 2019). Mouse models of Krabbe’s disease, another fatal leukodystrophy, showed similar microglial activation before myelin loss, with elevated innate immune markers even before microglial activation (Potter *et al.*, 2013; Snook *et al.*, 2014). The phenotype of one of the more prevalent leukodystrophies—Aicardi Goutières Syndrome (AGS)—mimics congenital cytomegalovirus brain infection and is classified as an interferonopathy to account for the upregulation of type I IFN-induced signal apparently central to pathology (Crow and Manel, 2015). Understanding whether the immune system is a driver of the disease rather a consequence may uncover new avenues for therapeutic interventions to treat these otherwise incurable diseases.

The development of animal models to study leukodystrophies *in vivo* has been invaluable for testing therapies (Priller *et al.*, 2001; Biffi *et al.*, 2004; Launay *et al.*, 2017; Marshall *et al.*, 2018), however many of those models fail to recapitulate the key aspects of the human disease. We have previously generated a zebrafish model for the RibonucleaseT2 (RNAseT2)-deficient leukoencephalopathy (Haud *et al.*, 2010). Similar to AGS, RNAseT2-deficient leukoencephalopathy is another severe leukodystrophy with clinical manifestations resembling a congenital cytomegalovirus brain infection. This disorder affects children in their first year of life and is characterised by white matter lesions, subcortical cysts and calcification, with severe psychomotor and sensorineural hearing impairments (Henneke *et al.*, 2009).

We and others have shown that loss of RNAseT2 causes a lysosomal storage disorder resulting in the accumulation of ribosomal RNA in neurons due to defects in ribosome autophagy (ribophagy) (Haud *et al.*, 2010; Hillwig *et al.*, 2011). Adult *rnaset2-deficient* zebrafish develop white matter lesions in adult brains, recapitulating this aspect of the human pathology. Furthermore *rnaset2* deficiency triggers increased expression of neuroinflammatory markers in the brain, further supporting a role for immune activation in driving pathogenesis in leukodystrophies (Haud *et al.*, 2010). However, as for most rare monogenic leukodystrophies, there is no treatment and the exact contribution of the immune system to the pathology is still obscure.

We therefore hypothesised that the early onset of RNAseT2-deficient leukoencephalopathy is caused by abnormal immune activity during brain development. In this current study, we utilise the *ex utero* development of zebrafish larvae and its optical transparency to study the role of RNAseT2 during early brain development. Using a combination of transgenic zebrafish and CRISPR/Cas9, we identified dysfunctional microglia in brains of *rnaset2-deficient* larvae displaying profound morphology defects and failing to clear apoptotic cells in the developing brain. This reduction in brain integrity was accompanied by locomotor defects at larval stages, and further validated in adults using behavioural tests. Furthermore, tissue-specific expression of *rnaset2* rescued microglial defects in *rnaset2* zebrafish mutants.

Our findings identify dysfunctional microglia as early markers of the pathology, highlighting a critical role for the immune system in disease development from its early stages. We propose a new mechanism by which failure of microglia to clear apoptotic neurons during development triggers irreversible neuroinflammation in the early brain, with negative impact on brain integrity. Our results identify the microglia as a new cellular target for therapies, opening the possibility of gene therapy and bone marrow transplant to treat this rare leukodystrophy.

## Results

### Transcriptome microarray profiling in *rnaset2^AO127^* mutant reveals increased immune responses in the brain

To investigate a role for the immune activation in *rnaset2*-deficient zebrafish, we adopted a unbiased approach to study differential gene expression in *rnaset2^AO127^* mutant. Microarray analysis was performed on whole *rnaset2^AO12^* and WT siblings during early development at 28 hours post fertilisation (hpf) and 3days post fertilisation (dpf), and on brain samples of 1-year-old adult zebrafish (Figure 1A). Biological pathway analysis identified 12 pathways differentially regulated in adult brain, with 7 involving the immune system (Figure 1B). The genes from ‘immune system’, ‘defence response’ and ‘innate immune response’ pathways were mostly upregulated in mutant samples (Figure 1B). 35% of the innate immune response genes are upregulated in mutant samples (Figure 1C), suggesting a strong immune response in the brain of mutants. Similar analysis did not reveal any significant pathway change in 28hpf samples, and only identified variations in cell cycle pathways at 3dpf (Sup. Figure 1). Additionally, our primary analysis showed that biological repeats from each time point clustered together, with the adult brain samples showing a clear separation from the embryonic time points (Sup. Figure 2). This suggests that the major dysregulations in gene expression happen in adulthood, with few differences being detectable in early embryonic timepoints looking at the whole animal level.

**Figure 1:**
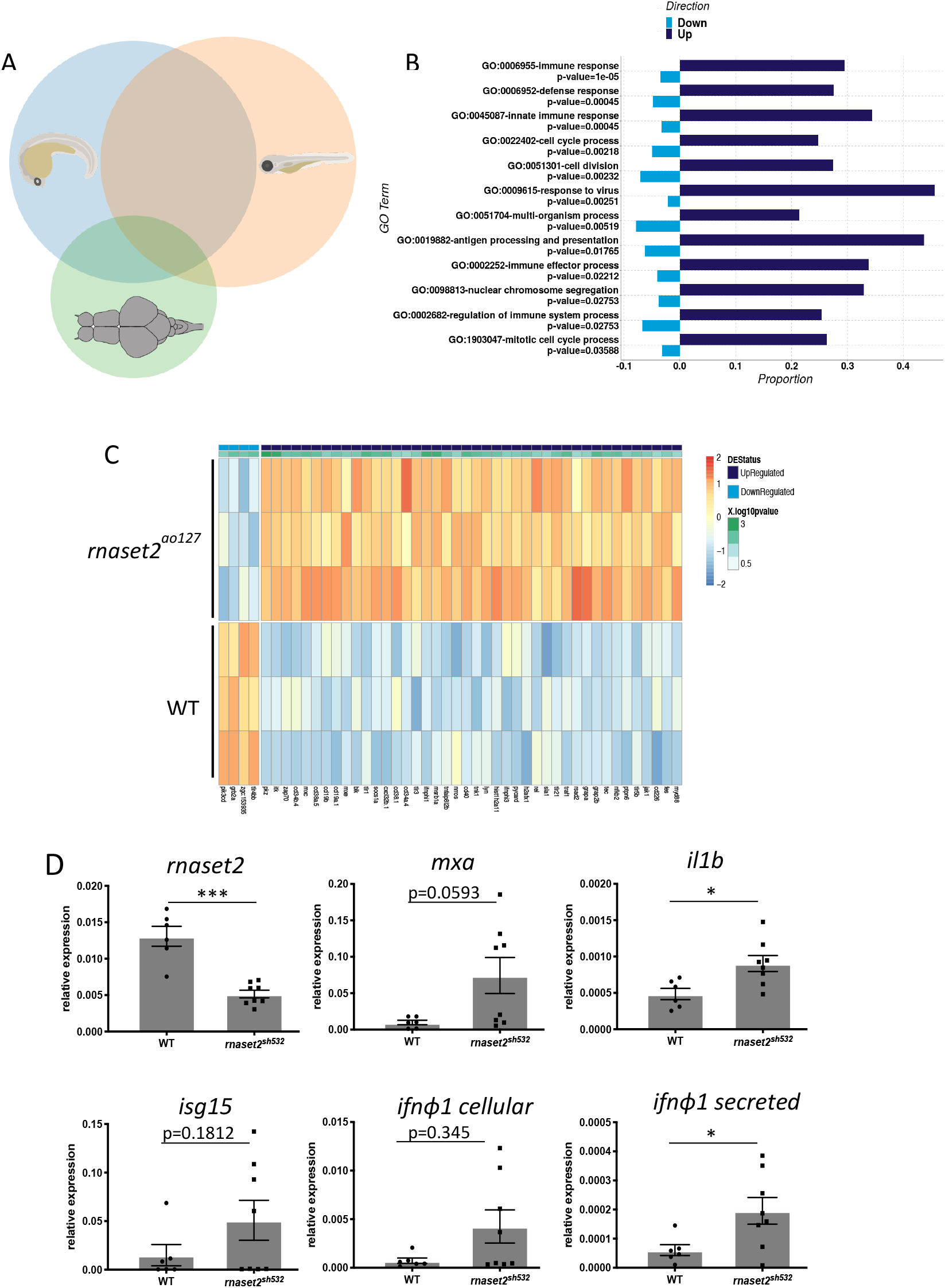
Immune system activity is upregulated in brains of *rnaset2* mutants. **A.** Venn diagram of differentially expressed genes across the three timepoints of *rnaset2^AO127^* mutant and WT samples used for microarray experiment. **B.** Differential expression (DE) graph showing the enriched pathways identified by clusterProfiler in adult brain. The proportion of individual genes belonging to each pathway is displayed in dark blue for up-regulated and light blue for down-regulated expression. The p-value for the enrichment of each pathway is stated below the pathway name. **C.** Heatmap of genes belonging to innate immune response GO pathway showing normalised expression values and all replicates of the adult time point. Genes are ordered according to the magnitude of difference between *rnaset2^AO127^* mutant and WT samples (with red indicating higher expression in mutant). Each row is annotated with the gene name and colour-coded bars at the top to indicate whether the gene is up-(dark blue) or down-(light blue) regulated, or not significant at the adjusted p-value threshold of 0.05. The −10log of the adjusted p-value is also shown. **D.** qPCR analysis of *rnaset2* and immune response genes: *mxa, il1beta, isg15, ifn Φ1 cellular* and *ifn Φ1 secreted* from dissected brain from 3-month-old WT siblings and *rnaset2^sh532^*. Expression relative to two reference genes combined *rpl13* and *ef1 a.* n=8 from 2 independent experiments, two-tailed Mann-Whitney U test, p value shown on individual graphs.

Interestingly, most genes in the GO ‘response to virus’ pathway were up-regulated in brains of adult *rnaset2* deficient zebrafish (Sup. Figure 3), providing supporting evidence that RNASET2-deficient leukodystrophy resembles a cytomegalovirus brain infection (Henneke *et al.*, 2009), and further demonstrating that our *rnaset2* mutants recapitulate the human disease.

### A CRISPR/Cas9 *rnaset2^sh532^* mutant recapitulates previous *rnaset2^AO127^* mutant

To further characterise the immune response triggered with the loss of rnaset2, we generated a second mutant allele by CRISPR/Cas9 in the *rnaset2* gene by targeting the same exon as our previous *rnaset2^AO127^* mutant, as our previous mutant was lost. This new *rnaset2^sh532^* mutant allele possesses an 8bp deletion in exon 5 (Sup. Figure 4A) which creates a STOP codon before the second catalytic domain resulting in a truncated rnaset2 protein (Sup. Figure 4B), similar to our previous mutant (Haud *et al.*, 2010). Homozygous and wild type (WT) sibling pairs were raised from a heterozygous in-cross and genotyped by high resolution melt curve analysis and/or restriction digest (Sup. Figure 4C, D). These fish were used to generate F2 homozygous *(rnaset2^sh532^*) and WT siblings which were viable and fertile, as observed with our previous mutant. The previous *rnaset2^AO127^* mutant had a characteristic high uptake of acridine orange, a dye accumulating in lysosomes highlighting the lysosomal storage defects (Haud *et al.*, 2010), a phenotype which is recapitulated by our *rnaset2^sh532^* mutant (Sup. Figure 5).

To determine whether immune response genes were also upregulated in the CRISPR mutant, we performed qPCR analysis on brains from 3-month-old *rnaset2^sh532^* mutant and WT siblings using upregulated genes identified in the microarray profiling of *rnaset2^AO127^*. *rnaset2* expression was decreased 3-fold compared to WT siblings (Figure 1D) suggesting nonsense-mediated RNA decay. Expression analysis of immune response genes confirmed upregulation of IL-1b *(il1beta),* interferon phi 1 genes *(ifnϕ1 cellular* and *ifnϕ1 secreted*) and antiviral genes (*mxa* and *isg15*) (Figure 1D). These results demonstrate that our new CRISPR/Cas9 *rnaset2^sh532^* mutant faithfully recapitulates the previous ENU-generated mutant.

### *rnaset2^sh532^* mutants exhibit abnormal behaviour in adult and larval stages

As the first generation of homozygous and WT siblings generated from genotyped F2 reached adulthood, we noticed a clear tilted swimming phenotype in 8-month-old *rnaset2^sh532^* mutants (Sup. Movie 1). Animals with tilted swimming behaviour were separated from the rest of the group for further analysis, after which this mild swimming phenotype developed into abnormal spiralling. They were therefore culled humanely before they deteriorated further. To perform an unbiased swimming behaviour analysis on the rest of the clutch, we used the ViewPoint system and recorded normal swimming behaviour for 10 minutes. WT adults exhibited exploratory behaviours, demonstrated by visiting all 4 quadrants of the tank equally (Figure 2A). Swimming and exploratory behaviours were significantly altered in *rnaset2^sh532^* adults, who swam in repetitive stereotyped patterns restricted to a single quadrant in which they spent the majority of their time (Figure 2B-C). Interestingly swimming speed and distance travelled was not different (Sup. Figure 6).

**Figure 2:**
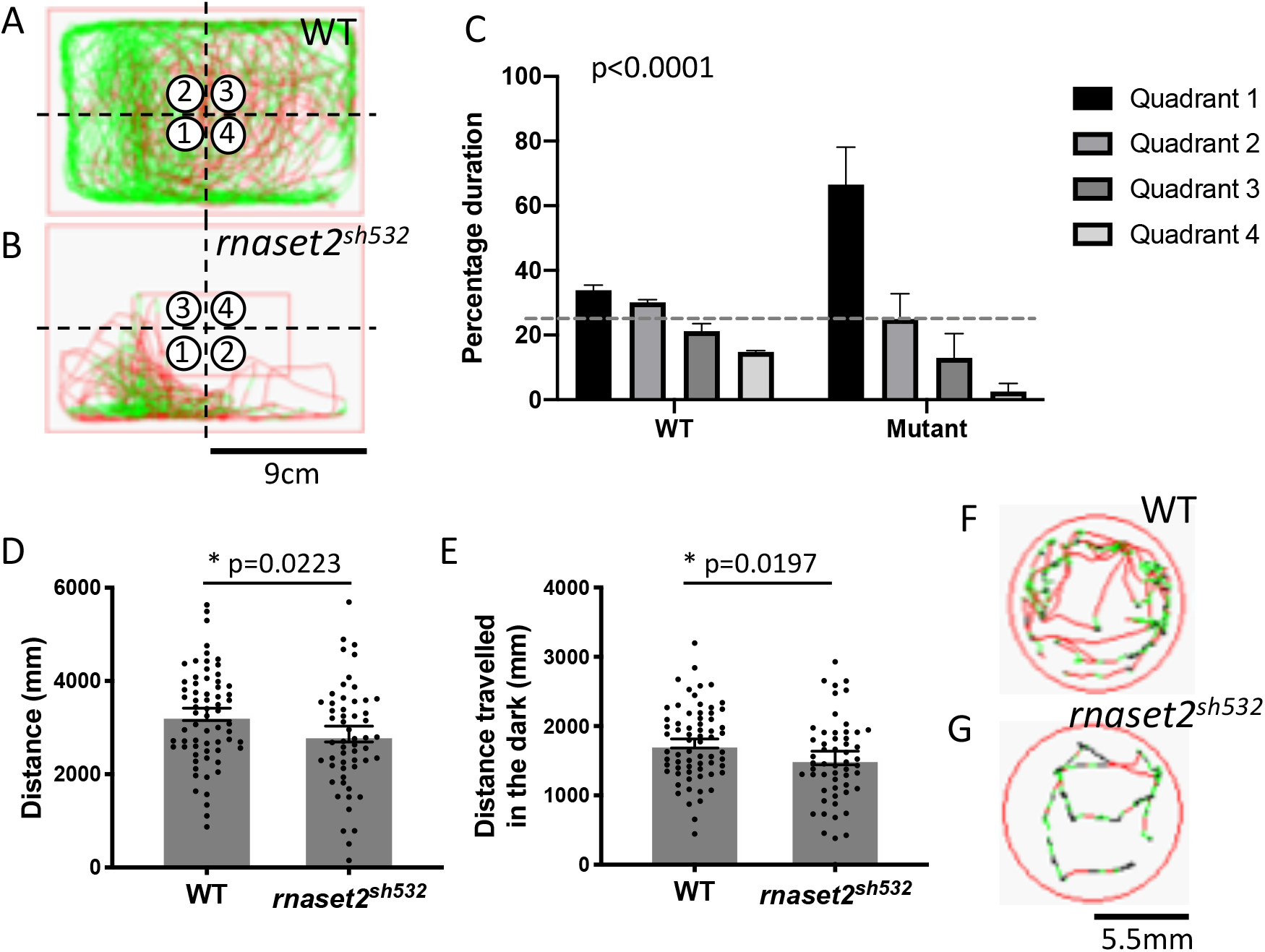
Locomotion tests reveals that *rnaset2*^sh532^ mutant have abnormal behaviour in adult and larval stages. **A, B.** Representative traces of movement from 8-month-old adults WT siblings (A) and *rnaset2^sh532^* (B) in all four quadrants with quadrant number 1 representing the quadrant most visited and quadrant 4 the least visited. **C.** Quantification of quadrant analysis in adult. n=4 using a contingency table and a Chi-square analysis pc0.0001. **D, E.** Swimming activity of 5dpf embryos as measured by distance travelled across a twenty minute period of alternating light and dark (D) and in dark phases only (E). n=56-67 larvae from 3 independent experiments, two-tailed Mann-Whitney U test p=0.0223 (D) and p=0.0482 (E). **F, G.** Examples of movement from 5dpf WT siblings (F) and *rnaset2^sh532^* larvae during 1 minute(G).

RNAseT2-deficient patients present with severe locomotor disabilities, including spasticity and dystonia, which arise in the first year of the patient life (Henneke *et al.*, 2009). To assess locomotion in early life, we performed locomotion tests at 5dpf using the Zebrabox ViewPoint system. We measured the swimming activity of 5dpf *rnaset2^sh532^* mutants relative to age-matched WT siblings over 20 minutes, alternating light and dark phases (Pant *et al.*, 2019). We found that *rnaset2^sh532^* larvae were hypoactive compared to WT siblings (Figure 2D-G). Total swimming distance over a period of twenty minutes of alternating light and dark was decreased in mutants compared to wild types, with hypoactivity accentuated in dark phases (Figure 2D, E p=0.0223, p=0.0482). These early onset of locomotor disabilities in the *rnaset2^sh532^* larvae suggest that the brain integrity is compromised during early stages of development.

### Increased numbers of apoptotic cells in 5dpf *rnaset2^sh532^* mutants are due to functionally impaired microglia

To characterise the early neuropathology and assess brain integrity in *rnaset2^sh532^* mutants, we analysed the level of neuronal apoptosis in the brain using the TUNEL ApopTag kit. A wave of neuronal apoptosis occurs in the developing zebrafish brain at 2dpf and is accompanied by an influx of embryonic macrophages to the brain (Herbomel *et al.*, 2001). These macrophages differentiate into microglia and begin to phagocyte apoptotic cells-a clearance that is completed by 5dpf (Casano *et al.*, 2016; Xu *et al.*, 2016). Quantification of apoptotic cell number in mutants identified a more than 2-fold increased at 5dpf (Figure 3A,C). Interestingly, we found no difference at the earlier timepoint of 3dpf (Sup. Figure 7). This shows that the initial number of dying neurons is unchanged in mutants and suggest a potential defect in apoptotic cell clearing by microglia.

**Figure 3:**
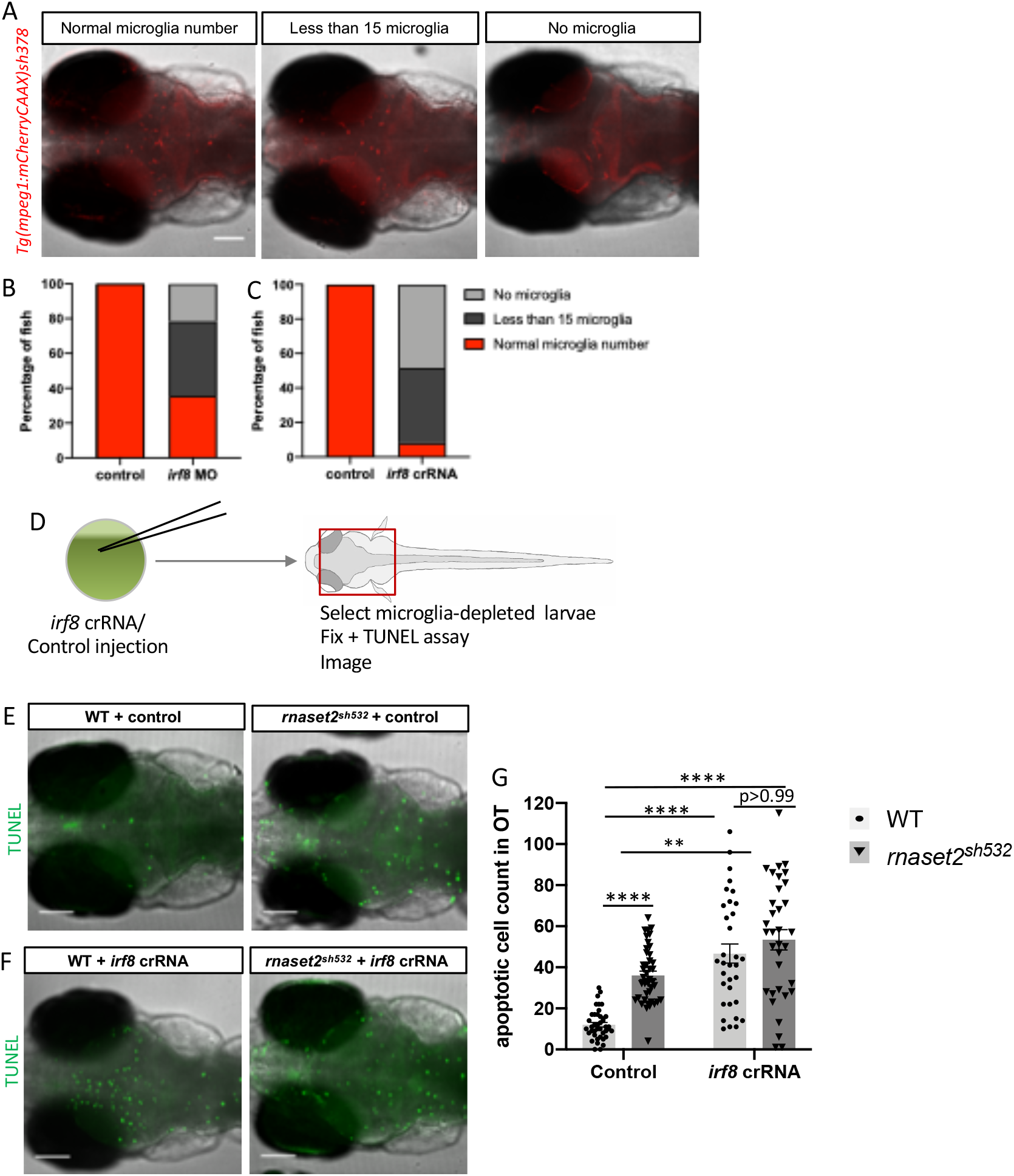
The increased number of apoptotic cells in 5dpf *rnaset2^sh532^* mutant is due to dysfunctional microglia. **A-C.** Injection of *irf8* crRNA is the optimal method for microglia depletion at 5dpf. A. Representative images of fish with normal microglia number, less than 15 and no microglia. Scale bar 50μm **B,C**. Quantification of microglia depletion after irf8 morpholino (MO) injection at 500μM and irf8 crRNA at 50μM. 3 biological replicates, n=120. **D.** Diagram of experimental flow. Injection of *irf8* crRNA or Control Tracr+Cas9 was performed into 1-cell stage embryos, which were raised until 5dpf and microglia-depleted larvae were selected in *irf8* crRNA injected animals. Control and depleted larvae were fixed and stained for apoptotic cells using TUNEL assay. **E, F.** Representative images of apoptotic cells (as visualised by TUNEL staining) in brains of 5dpf *rnaset2^sh532^* mutants and WT siblings in Cas9+TracrRNA injected control (E) and following injection of *irf8* crRNA (F). **G.** Quantification of apoptotic cells in the optic tectum (OT). n=125-131 from 3 independent experiments, using multiple two-tailed U-test with Bonferroni’s multiple comparisons test (***: p<0.001, ****; p<0.0001).

To confirm that this increased in number of apoptotic cells in mutants are due to microglial dysfunction and not to increased dying neurons, we depleted macrophages and microglia by targeting *irf8* using the CRISPR/Cas9 system. Using the CRISPR/Cas9 approach resulted in a more robust depletion of microglia at 5dpf compared to the *irf8* antisense morpholino (Figure 3 A-C), an approach commonly used to deplete macrophages and microglia at earlier stages (Li *et al.*, 2011; Shiau *et al.*, 2015). As expected and previously shown (Villani *et al.*, 2019), depletion of microglia in WT resulted in a 3-fold increase numbers of apoptotic cells compared to control injected larvae at 5dpf (Figure 3D,E,G). Depletion of microglia in *rnaset2^sh532^* mutants resulted in the same number of apoptotic cells than in depleted WT (Figure 3D,F,G). We therefore concluded that the rate of neuronal apoptosis is normal in mutants and that the increased number of apoptotic cells in mutant is due to a defect in clearing of dead cells by microglia.

### Microglia display impaired morphology and increased number in *rnaset2^sh532^* mutants

To further characterise microglial defects in *rnaset2^sh532^* mutants, we analysed microglial number and morphology using the fluorescent reporter line *Tg(mpeg1:mCherryCAAX)sh378* labelling the membrane of macrophages and microglia. Confocal images of the optic tectum at 5dpf revealed that microglia were more numerous in mutant brains at 5dpf (Figure 4A-C, p= 0.0077). Subsequent morphology analysis identified that microglia in mutant brains were more circular with a higher circularity index (Figure 4A,D,E, p<0.0001) and contained a high number of vacuoles labelled using the membrane transgenic line *Tg(mpeg1:mCherryCAAX)sh378* (Sup. Figure 8). This circular and engorged phenotype was reduced in homozygous mutants generated from a heterozygous in-cross, suggesting that maternal contribution of *rnaset2* in early development is critical for maintaining a normal microglia morphology (Sup. Figure 9). To determine if loss of *rnaset2* affects all macrophages, we assessed macrophage innate immune function by studying responses to tissue damage induced by caudal tail fin transection (Morales and Allende, 2019). No difference was identified in macrophage recruitment to the wound site at 24hpi (Figure 4A,F,G), suggesting that loss of *rnaset2* disrupts microglial function in the CNS without affecting the innate immune responses in the periphery.

**Figure 4:**
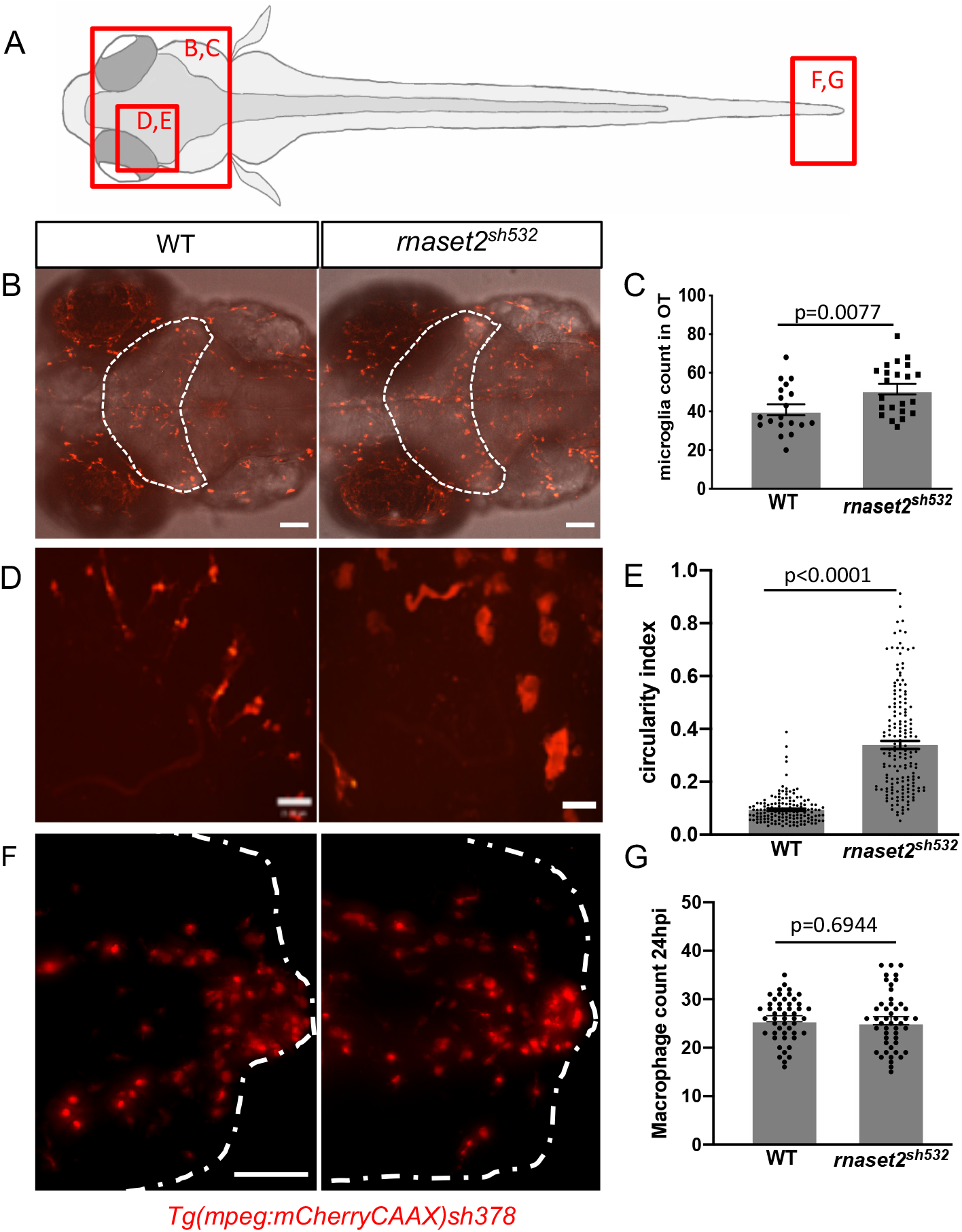
Microglia display impaired morphology and increased number in *rnaset2^sh532^* mutant. **A.** Diagram of a dorsal view of 5dpf zebrafish larvae with red boxes highlighting the brain and tail fin regions imaged in the next panels. **B, C.** Microglia count in the optic tectum of 5dpf WT siblings and *rnaset2^sh532^* larvae. B. Representative images of the whole head of 5dpf larvae with optic tectum outlined (dotted white line). Scale bar 70μm. **C.** Quantification of microglia number in the optic tectum. n=154-173 larvae from 3 independent experiments, two-tailed Mann-Whitney U test p<0.0001. **D, E.** Analysis of microglia circularity in the optic tectum in 5dpf WT siblings and *rnaset2^sh532^* larvae. **D.** Representative images of microglia morphology using the 40x objective. Scale bar 21μm E. Quantification of microglia circularity using the circularity index analysis from Fiji. n=48-50 larvae from 3 independent experiments, two-tailed Mann-Whitney U test p=0.6944. **F,G.** Analysis of macrophage numbers at tissue damage site in 3dpf WT siblings and *rnaset2^sh532^* larvae. **F.** Side view of tail fin injury representative images of macrophages at wound site at 24 hours post injury. Scale bar 50μm G. Quantification of macrophage numbers at the wound site 24 hours post injury. n=48-50 larvae from 3 independent experiments, two-tailed Mann-Whitney U test p=0.6944.

### Tissue-specific rescue of *rnaset2* restores microglial morphology in *rnaset2^sh532^* mutant

To further investigate the microglial phenotype identified in *rnaset2^sh532^* mutants, we rescued *rnaset2* expression by re-expressing WT *rnaset2* in the *rnaset2^sh532^* mutant. Rescue plasmids were built using gateway cloning to rescue expression in whole larvae (using the ubiquitous *ubi* promoter), specifically in macrophages/microglia (using the *mpeg1* promoter), or specifically in neurons (using the *huc* promoter). All constructs contained a WT *rnaset2* middle entry vector, and a blue eye marker to allow screening of positive larvae. Ubiquitous expression of *rnaset2* fully rescued the brain phenotype observed in the *rnaset2^sh532^* mutants, such that there was no difference in microglia circularity index compared to WT microglia (Figure 5A,B,D,E, p<0.0001). Re-expression of *rnaset2* specifically in macrophages and microglia using the *mpeg:rnaset2* construct also rescued the circularity index, but to a lesser degree than in WT (Figure 5A,B,D, p<0.001, Sup. Figure 10A). Interestingly, rescue of *rnaset2* specifically in neurons using the *huc:rnaset2* construct achieved complete rescue of the microglia circular phenotype as seen when using the ubiquitous rescue (Figure 5A,B,E, p<0.0001, Sup. Figure 10B). These results demonstrate that rescuing rnaset2 specifically in microglia or in neurons can alleviate developmental microglial defects. This highlights cell-specific strategies for targeted therapy, representing exciting therapeutic avenues for treatment of *rnaset2^sh532^* deficient leukoencephalopathy.

**Figure 5:**
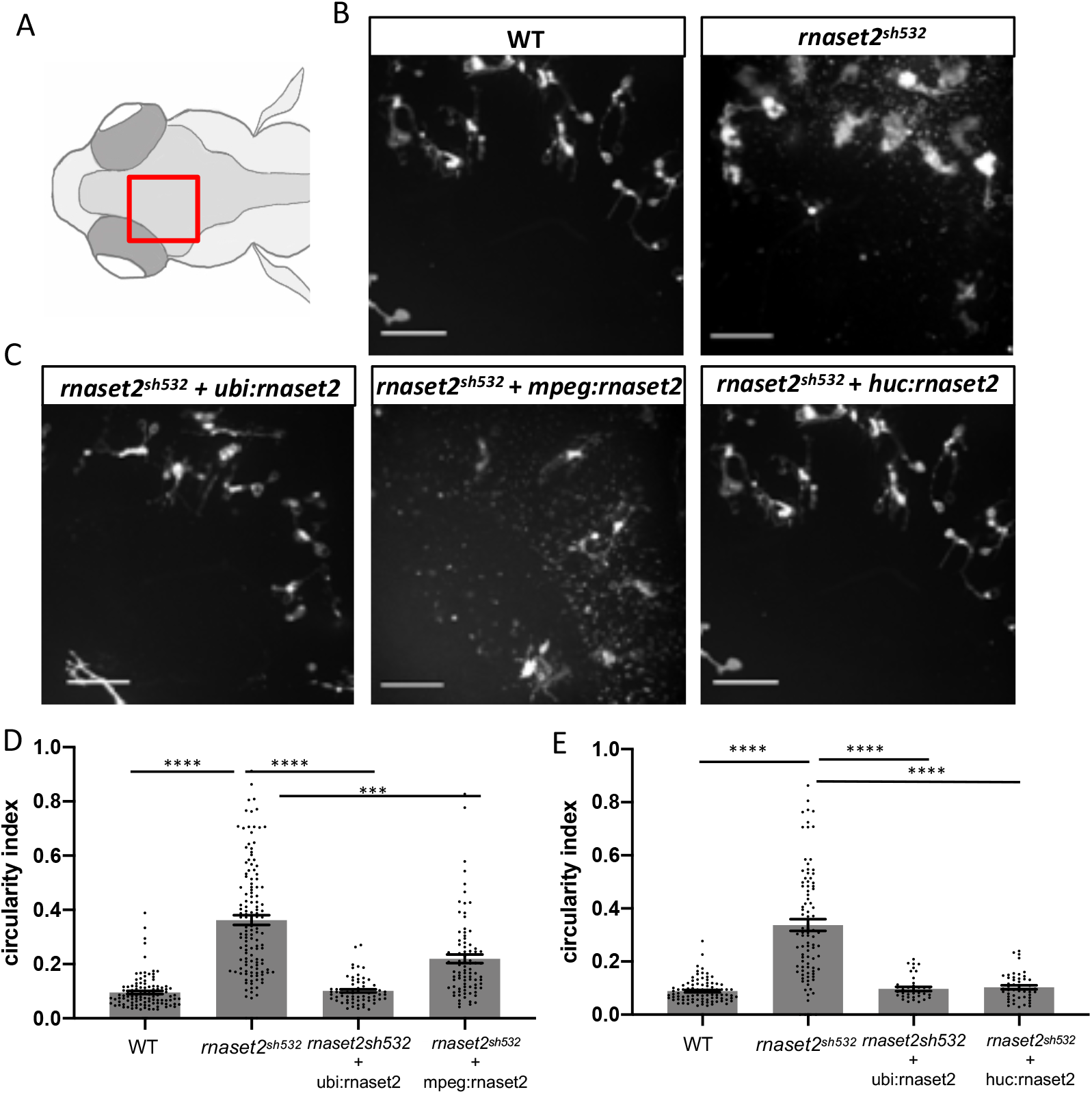
Tissue-specific rescue of *rnaset2* restores microglial morphology in *rnaset2^sh532^* mutant. **A.** Diagram of a dorsal view of a 5dpf zebrafish head with red box highlighting the region imaged in the next panels. **B.** Representative images of microglia morphology in 5dpf WT siblings and *rnaset2^sh532^* larvae using the 40x objective. Scale bar 100μm **C.** Representative images of microglia morphology using the 40x objective in 5dpf *rnaset2^sh532^* larvae injected at one-cell stage to rescue expression of the *rnaset2* gene either ubiquitously, or in macrophages and microglia, or in neurons. Scale bar 100μm. **D. E.** Quantification of microglia circularity after rescue of *rnaset2* expression ubiquitously and in macrophages/microglia **(D)** and after rescue of *rnaset2* expression ubiquitously and in neurons **(E)** using the circularity index analysis from Fiji. n=64-132 larvae from 3 independent experiments, Kruskal-Wallis test with Dunn’s multiple comparisons p<0.0001.

## Discussion

We have identified dysfunctional microglia as a new early marker of the pathology in RNAseT2-deficient leukoencephalopathy. Our results also demonstrate that microglia-specific interventions can alleviate microglial defects in *rnaset2*-deficient zebrafish, therefore highlighting a cell specific therapeutic approach to be investigated in patients.

RNAseT2-deficient leukoencephalopathy was already proposed to be classified as a lysosomal storage disorder (LSD) from our previous study, showing accumulation of ribosomal RNA (rRNA) in neurons (Haud *et al.*, 2010). The specific rounded and engorged microglial phenotype in *rnaset2^sh532^* mutants is indeed reminiscent of an LSD and reinforces our initial proposal. Although these microglial defects have not yet been described in RNAseT2-deficient leukoencephalopathy patients, they still represent an early marker of the pathology and a potential therapeutic target for RNASET2-deficient leukoencephalopathy.

In our model, we showed increased number of apoptotic bodies in brain tissue caused by failure of microglia to clear dying cells. Defects in the clearance of apoptotic neurons by microglia can promote additional neuronal death (Peri and Nüsslein-Volhard, 2008; Casano *et al.*, 2016; Villani *et al.*, 2019) and have been associated with WMLs in neurodegenerative conditions and ageing (Raj *et al.*, 2017; Rissanen *et al.*, 2018). Additionally, increased apoptosis in the brain acts as a signal for brain entry of macrophages (Casano *et al.*, 2016; Xu *et al.*, 2016), which can explain the increased number of microglia observed in mutant brains. Cumulative arrival of microglia in the brain could eventually clear up the brain from apoptotic bodies, but at the expense of accumulating engorged microglia which could potentially secrete stress factor and become neurotoxic. This could trigger a ‘snowball’ effect, by which *rnaset2*-deficient microglia phagocytose a fixed number of dying neurons before reaching saturation-resulting in more microglia entering the brain to create a constant neuroinflammatory state. Future microglia-specific transcriptomic studies will be informative to elucidate their inflammatory profile.

Tissue-specific rescue of *rnaset2* expression using a neuronal promoter fully restored microglia morphology compared to a partial rescue with a microglia promoter. This could be due to the sheer number of neurons expressing *rnaset2* in the case of the neuronal rescue, as opposed to a few microglia only in the case of the use of the microglia promoter. Deficient microglia could inherit the corrected version of the protein by engulfing rescued neuron. Cross-correction, the phenomenon at the origin of enzyme replacement therapy, can also be a mechanism as neurons could be secreting extracellularly the rnaset2 protein which could then be taken up by surrounding microglia (Fratantoni *et al.*, 1968).

Most importantly, to our knowledge, our zebrafish model is the first man-made animal model of a leukodystrophy that fully recapitulate the patient brain phenotype throughout development and upon adulthood. Murine models of leukodystrophies have failed to reproduce all aspect of the human disease with a recent RNASET2-deficient rat model showed deficiency in object recognition memory but showed no locomotor or spatial memory defects (Sinkevicius *et al.*, 2018). The *rnaset2* zebrafish mutants shows hypoactivity in larval stages and tilted swimming in adulthood with an impaired exploratory phenotype. The adult tilted swimming phenotype can be caused by defects in the vestibular righting reflex or motor deficit (Kalueff *et al.*, 2013). Vestibular defects a due to impaired ear development or impairment of sensory neurons within the inner ear (Whitfield, 2002) and recapitulates the sensorineural hearing loss diagnosed in patients (Henneke *et al.*, 2009). Therefore, our study places the *rnaset2*^sh532^ zebrafish mutant at the forefront of leukodystrophy pre-clinical animal models and will be used to develop therapies targeting the microglial population.

Our study identifies an important role for the ribonuclease T2 protein in early brain development and reveals that deficient microglia could underpin the pathology of RNAseT2-deficient leukoencephalopathy. Here we report multiple features of neuroinflammation including apoptosis, increased immune response, white matter pathology and neuromotor deficits in the form of impaired locomotion in larval and adult *rnaset2*-deficient zebrafish mutants. We provide evidence to focus on therapies that can target the microglial population, such as gene therapy and haematopoietic stem cell transplantation. These therapies are already being used in clinical trials for LSDs and other types of leukodystrophies (Penati *et al.*, 2017; Ohashi, 2019; Schiller *et al.*, 2019) and could therefore be adapted to treat children suffering from RNAseT2-deficient leukoencephalopathy.

## Materials and Methods

### Zebrafish husbandry and ethics

All zebrafish were raised in the Bateson Centre at the University of Sheffield in UK Home Office approved aquaria and maintained following standard protocols (Nüsslein-Volhard and Dham, 2002). Tanks were maintained at 28°C with a continuous re-circulating water supply and a daily light/dark cycle of 14/10 hours. All procedures were performed on embryos less than 5.2 dpf which were therefore outside of the Animals (Scientific Procedures) Act, to standards set by the UK Home Office. We used the *Tg(mpeg1:mCherryCAAX)sh378* labelling the membrane of macrophages and microglia (Bojarczuk *et al.*, 2016), the previously ENU-generated *rnaset2^AO127^* mutant and *nacre* wild type (WT) to generate the new CRISPR/Cas9 *rnaset2^sh532^* mutant.

### Microarray experiment and analysis

RNA was isolated from pooled dechorionated embryos at 28 hour post fertilisation (hpf), 3 day post fertilisation (hpf) and dissected whole adult brains using TRIzol (Invitrogen), following the manufacturer’s protocol. The integrity of the RNA was confirmed using the Agilent 2100 BioAnalyzer, using only samples with an RNA integrity number of at least 8.

Genome-wide expression profiling was performed using the Zebrafish (v3) Gene Expression 4×44K Microarray (Agilent G2519F) containing 43803 probes according to the manufacturer’s instructions. The probe sequences for this array platform were re-annotated using the ReAnnotator pipeline (Arloth *et al.*, 2015) against reference genome danRer11 in order to obtain updated gene symbols.

Scanned raw data were normalised using quantile normalisation, and then filtered using the ReAnnotator results so that only probes matching the exons of their target gene were retained. In the case of genes with more than 1 probe, the probe with the highest inter-quartile range was chosen as the representative probe for the gene.

Differential expression analysis was conducted using a linear model framework and empirical Bayes’ shrinkage implemented in the limma Bioconductor package (Ritchie *et al.*, 2015). Genes with an adjusted p-value <0.05 were taken forward to enrichment analysis with the clusterProfiler Bioconductor package (Yu *et al.*, 2012).

### CRISPR/Cas9 design and injection

Synthetic SygRNA^®^ consisting of gene specific CRISPR RNAs (crRNA) (Sigma) and transactivating RNAs (tracrRNA) (Merck) in combination with cas9 nuclease protein (Merck) was used for gene editing. TracrRNA and crRNA were resuspended to a concentration of 50μM in nuclease free water containing 10mM Tris-HCl ph8. SygRNA^®^ complexes were assembled on ice immediately before injection using a 1:1:1 ratio of crRNA:tracrRNA:Cas9 protein. We used the following crRNA sequences, where the PAM site is indicated in brackets: *rnaset2* (CCG)AGATCTGCTAGAACCATCTT, *irf8* GCGGTCGCAGACTGAAACAG(TGG). A 2nl drop of SygRNA^®^:Cas9 protein complex was injected into one-cell stage embryos. *rnaset2* crispants were raised and screened (see below) to select a suitable mutation for a stable line.

Successful *irf8* crispant were phenotypically identified by injecting the SygRNA^®^:Cas9 complex into the *Tg(mpeg1:mCherryCAAX)sh378* and selecting embryos with no microglia at 5dpf. Control injection contained TracrRNA and Cas9 nuclease only.

### Generation of *rnaset^sh532^* mutant

To determine the efficiency of CRISPR/Cas9 to induce site-specific mutations in injected larvae, we used high-resolution melt curve (HMRC) analysis or PCR followed by *Mwo1* restriction digest (Sup. figure 4C,D). Genomic DNA (gDNA) was extracted from individual larvae at 2dpf. Larvae were placed individually in 90μl 50mM NaOH and boiled at 95° for 20 minutes. 10μl Tris-HCL ph8 was added to adjust the pH and mixed thoroughly. Gene specific primers were designed using the Primer 3 web tool (http://primer3.ut.ee/) and primers with the following sequences were used to amplify a 142bp region: HRMCrnaset2_fw ACATACTACCAGAAATGGAG, HMRCrnaset2_rev GTAGTGCCTAAATGCATTTG. HMRC analysis was run using the CFX96 Bio-rad qPCR machine with the Bio-Rad Precision Melt Analysis software using the following reaction: gDNA 1 μl, 5ul DyNAmo Flash SYBR Green (Thermofisher), 0.5μl of each primers 10μM and 4μl of water. The program used was: Step 1: 95°C for 2min, Step 2: 95°C for 10sec, Step 3: 60°C for 30sec, Step 4: 72°C for 30sec, Step 5: 95°C for 30sec, Step 6: 60°C for 10min, Step 7: 95°C for 20sec with Step 2 to 4 repeated 44 times and increment of 0.2°C every 10sec between Step 6 and Step 7.

Potential founders were outcrossed and gDNA was extracted from 8 embryos from each progeny. HMRC was used to identify INDELS in progenies from founder 2 and 5, which were subsequently raised. Individual F1 adults were fin clipped and DNA of individuals with INDELS was sequenced using same genotyping primers as above. An 8 base pair deletion causing a frame shift and an early stop codon in a similar region as our previous *rnaset2* mutant was kept, given the ‘*sh532*’ code and outcrossed to the *Tg(mpeg1:mCherryCAAX)sh378* line. Heterozygous animals were in-crossed and genotyped. Homozygous (*rnaset2*^sh532^) and WT siblings (WT) were selected, separated and raised to adulthood. All experiments were done on in-crosses from each tank, except when stated otherwise in Sup. Figure 9.

### Adult behaviour tests

As above, behavioural analysis was performed using the Zebrabox tracking system and Zebralab software (ViewPoint Life Science, France). 8- and 3-month-old wild type and mutant fish were matched for sex and size, before being placed individually into open field tanks and allowed to habituate for 30 minutes before recording. The walls of each tank were covered with white paper to ensure the animals were unable to interact or be distracted by their surroundings. Movement was tracked over a period of 10 minutes under constant light conditions. For quadrant analysis, the Zebrabox software was programmed such that the open field tank was divided into four equal quadrants, where quadrant 1 was the most active quadrant while quadrant 4 was the least. Distance travelled by each fish was pooled and averaged—both in total, and within each quadrant—in Microsoft Excel, with the resulting data analysed in GraphPad Prism.

### Larval behaviour light/dark behaviour test

Behavioural analysis was performed using the Zebrabox tracking system and Zebralab software (ViewPoint Life Science, France).

To assess swimming behaviour in response to light, 5dpf embryos were transferred into 48-well plates (one embryo per well) and allowed to habituate overnight. Swimming distance during a one-minute interval was recorded for twenty minutes during an alternating dark-light protocol (five minutes dark, followed by five minutes light). Total distance was calculated as the sum of the distance swam across all twenty intervals for a single fish. Response to changes in light intensity was defined as the distance swam in the minute after each light-to-dark or dark-to-light transition. Distance travelled by each fish was pooled and averaged and the resulting data was analysed using GraphPad Prism. Outliers that had not been tracked accurately over the recording period were manually excluded based on abnormal angles and straight lines generated by the Zebralab software.

### *irf8* Morpholino knockdown

The *irf8* modified antisense oligonucleotide-morpholino (Gene Tools) was used as previously reported by injecting 1nl of 0.5mM irf8 morpholino in the yolk of 1-cell stage embryos (Li *et al.*, 2011).

### Acridine Orange assay

A pool of twenty 5dpf WT and *rnaset2^sh532^* mutant zebrafish larvae were incubated at 27°C for 20 minutes in 5μg/ml of Acridine Orange (Life Technologies) diluted in zebrafish E3 medium. The larvae were rinsed three times for 10 minutes in clean E3. Larvae were then sedated and chopped finely with a scalpel then transferred to a tube containing PBS with Liberase (Roche, reference 05401020001) at 40μg/ml. Samples were incubated at 37°C for 30 minutes, with triturating of the mixture every 10 minutes to dissociate into a single cell suspension. Samples were centrifuged at 1000g for 6 minutes and resuspended in Leibovitz 15 media containing 20% FBS and 5mM EDTA. After filtration to remove clumps of cells, samples were analysed using an ATTUNE Flow cytometry machine using the blue laser.

### Quantitative PCR

RNA from brains of 3-month-old zebrafish was extracted in 500μl of Trizol (Invitrogen) and purified by adding 100μl of chloroform. After vigorous shaking, samples were spun down at 4°C for 20 minutes and the top phase was transferred to a new tube containing 300μl of isopropanol to precipitate the RNA overnight at −20°C. RNA was washed in 70% ethanol and dried before being resuspended in RNAse-free water. cDNA synthesis was performed using 2μg of RNA and the SuperScript II kit (Invitrogen) following manufacturer instructions. cDNA was diluted in 1:20 for qPCR. qPCR primers (Supplementary Table 1) were tested for efficiency (85%-105%) using a cDNA serial dilution. The qPCR reaction was run in a CFX96 Bio-Rad machine as follow: 2μl of cDNA, 5ul DyNAmo Flash SYBR Green (Thermofisher), 0.5μl of each primers 10μM and 3μl of water. The program used was: Step 1: 95°C for 2min, Step 2: 95°C for 10sec, Step 3: 60°C for 30sec, Step 4: 72°C for 25sec, Step 5: 95°C for 30sec, Step 6: 65°C for 10sec, Step 7: 95°C for 20sec with Step 2 to 4 repeated 39 times and increment of 0.2°C every 10sec between Step 6 and Step 7.

ΔCT was calculated using combined *rpl13* and *ef1 α* as reference genes and expression relative to endogenous control was calculated using the 2^^^(-ΔCT).

### Microglia morphology and count

To assess microglia morphology, 5dpf WT and *rnaset2^sh532^* mutant zebrafish larvae were anaesthetised and imbedded in low melting point agarose containing tricaine (0.168 mg/ml; Sigma-Aldrich) and imaged using a 40x objective on a UltraVIEW VoX spinning disk confocal microscope (PerkinElmer Life and Analytical Sciences) confocal spinning disk microscope. 100micron stacks were acquired using 0.5μm slices. For image analysis, stacks of 50μm were used to create a maximum projection and contours of each microglia (avoiding pigment cells) were drawn using a pen tablet (Intuos from Wacom). Using Fiji, we measured the circularity index (0-1) of each microglia and used these values to assess the circularity of each cell with the value of 0 being not circular and 1 as being perfectly circular.

The same images were used to quantify the number of vacuoles in each microglia. We used the labelled microglia membrane from *Tg(mpeg1:mCherryCAAX)sh378* to manually count vacuoles. Number of vacuoles in microglia exceeding 20 were capped due to the difficulty to be accurate after this number.

For microglia count, 5dpf larvae were imaged under the 10x objective of the spinning disk microscope. Stacks of 100μm were generated with 2μm thickness and microglia were counted in the optic tectum region of each larvae by drawing a region of interest around the optic tectum first using the reference brightfield image.

### Macrophage recruitment assays

To induce an inflammatory response, 2dpf WT and *rnaset2^sh532^* mutant zebrafish larvae in the *Tg(mpeg:mCherryCAAX)sh378* background were dechorionated and anaesthetised using Tricaine. Tail-fins were transected consistently using a scalpel blade (5mm depth, WPI) by slicing immediately posterior to the circulatory loop, ensuring the circulatory loop remained intact as previously described. Larvae were left to recover in fresh E3 media at 28°C. Larvae were mounted for imaging 24 hours post injury in a 1% low melting point agarose solution (Sigma-Aldrich) containing tricaine. Images were taken using an Andor Zyla 5 camera (Nikon) and an mCherry specific filter with excitation at 561nm. 10 z-planes were captured per larvae over a focal range of 100μm. Maximum intensity projections were generated by NIS elements (Nikon) to visualise all 10 z-planes.

### TUNEL assay

Fish larvae at 5dpf were fixed in 4% PFA and the TUNEL assay was performed according to standard protocol using ApopTag Kit (Millipore). Briefly, 5dpf larvae were incubated with proteinase K (20μg/ml) for 2 hours to permeabilise the tissue. Samples were fixed for 20 minutes in PFA at room temperature before being placed at −20°C in 1:2 acetone: ethanol for 7 minutes. Following incubation at 37°C with 50μl equilibration buffer for 1 hour, the reaction solution (16μl TdT enzyme and 30μl reaction buffer; Apoptag Kit) was added to the embryos— again, incubating at 37°C for 90 minutes. Embryos were then placed in 200μl stop buffer (Apoptag kit) for 2 hours at 37°C, before placing in antibody and blocking solution (62μl anti-Dig Fluorescein and 68μl blocking solution) overnight at 4°C. The following morning, samples were thoroughly washed in PBST (PST + 0.1% Tween) and fixed in PFA for 30 minutes at room temperature. All liquid was removed and embryos thoroughly rinsed with PBST between each stage (5 minute washes at room temperature, repeated three to four times as needed). Following completion of TUNEL staining, samples were imaged on the inverted UltraVIEW VoX spinning disk confocal microscope (PerkinElmer Life and Analytical Sciences) using the brightfield and GFP channel by acquiring stacks of approximately 100μm with 2μm per slice. Using Fiji, the optic tectum region was outlined with the free hand drawing tool using the brightfield image and counting of number of apoptotic cells was performed using the automatic thresholding and particle count. Counting was performed for all samples from each experiment within the same day to minimise intra-observer variation.

### Statistical analysis

All statistical analysis were performed in GraphPad Prism where data was entered using either a column (2 samples, 1 variable only) or a grouped table (more than 2 samples or 2 variables). Sample distribution was assessed using frequency of distribution analysis. Non-parametric tests were used for not normally distributed dataset with post-test (for multiple comparisons) described for each experiment. All experiments were repeated at least twice, with the number of biological replicates and n (experimental unit) number stated for each experiment in figure legends. Contingency tables were used for adult zebrafish behaviour analysis with Fisher’s exact-test.

p values are indicated and a star system is used instead for graph with multiple comparisons: *=p<0.05, **=p<0.01, ***=p<0.001, ****=p<0.0001. Following the recommendation of the American Statistical Association we do not associate a specific p value with significance (Wasserstein *et al.*, 2019).

### Cloning of rescue constructs

The *rnaset2* cDNA was amplified from the PCGlobin vector previously used (Haud *et al.*, 2010) using the following *Attb* primers: rnaset2_AttB1fwd GGGGACAAGTTTGTACAAAAAAGCAGG-CTGGATGAGATTCATTGCATTTGCTG, rnaset2_AttB2rev GGGGACCACTTTGTACAAGAA-AGCTGGGTGCTACGCTTGCACCGGTGGGTA. Using the BP reaction, the PCR product was cloned into the p221DONOR vector which was used for the following LR reactions. To create the positive control construct *ubi:rnaset2:pA* construct, we used the p5’E-ubi (Mosimann *et al.*, 2011), pME-*rnaset2*, p3’E-polyA into the *cryCFPpDest* vector. To create the *mpeg1:rnaset2:pA* construct, we used the p5’E-*mpeg1* (Ellett *et al.*, 2011), pME-*rnaset2*, p3’E-polyA into the *cryCFPpDest* vector. To create the *huc:rnaset2:pA* construct, we used the p5’E-*huc* (cloned into p5’E using the following primers and restriction sites: Xhohuc_fwd CGACTGCTCGAGCTTCCGGCTCGTATGTTGTG and Bamhuc_rev GCAGGATCCGGTCCTTCGATTTGCAGGTC), pME-*rnaset2*, p3’E-polyA into the *cryCFPpDest* vector. Blue-eyed larvae were selected at 5dpf before imaging.

## Acknowledgements

We thank David Drew for technical support, the staff of the light microscopy facility Dr Darren Robinson and Nick Van Hateren and the staff of the Sheffield Zebrafish Screening Unit Jean-Paul Ashton and Sarah Baxendale. We thank the Bateson Centre aquaria staff for their assistance with zebrafish husbandry. We thank Dr Daniel Lysko, Dr Will Talbot and Dr Ryan McDonald for sharing the *irf8* crRNA sequence and Dr Stone Elworthy for the 5’E-*huc* vector. We thank Dr Simon Johnston, Dr Katy Henry and Dr Andy Grierson for helpful feedback on our manuscript.

## Author contributions

Conceptualization, NH and SAR; Methodology, NH and TW; Investigation, NH, HAR, HMI, JP, MD; Writing – Original Draft, NH and HR; Writing – Review & Editing, NH, HMI and SAR; Funding Acquisition, NH and SAR; Visualization, NH and HR; Resources and Funding Acquisition, NH, SAR, JG and MH; Supervision, NH and SAR.

## Supplementary files

**Supplementary Figure 1:**
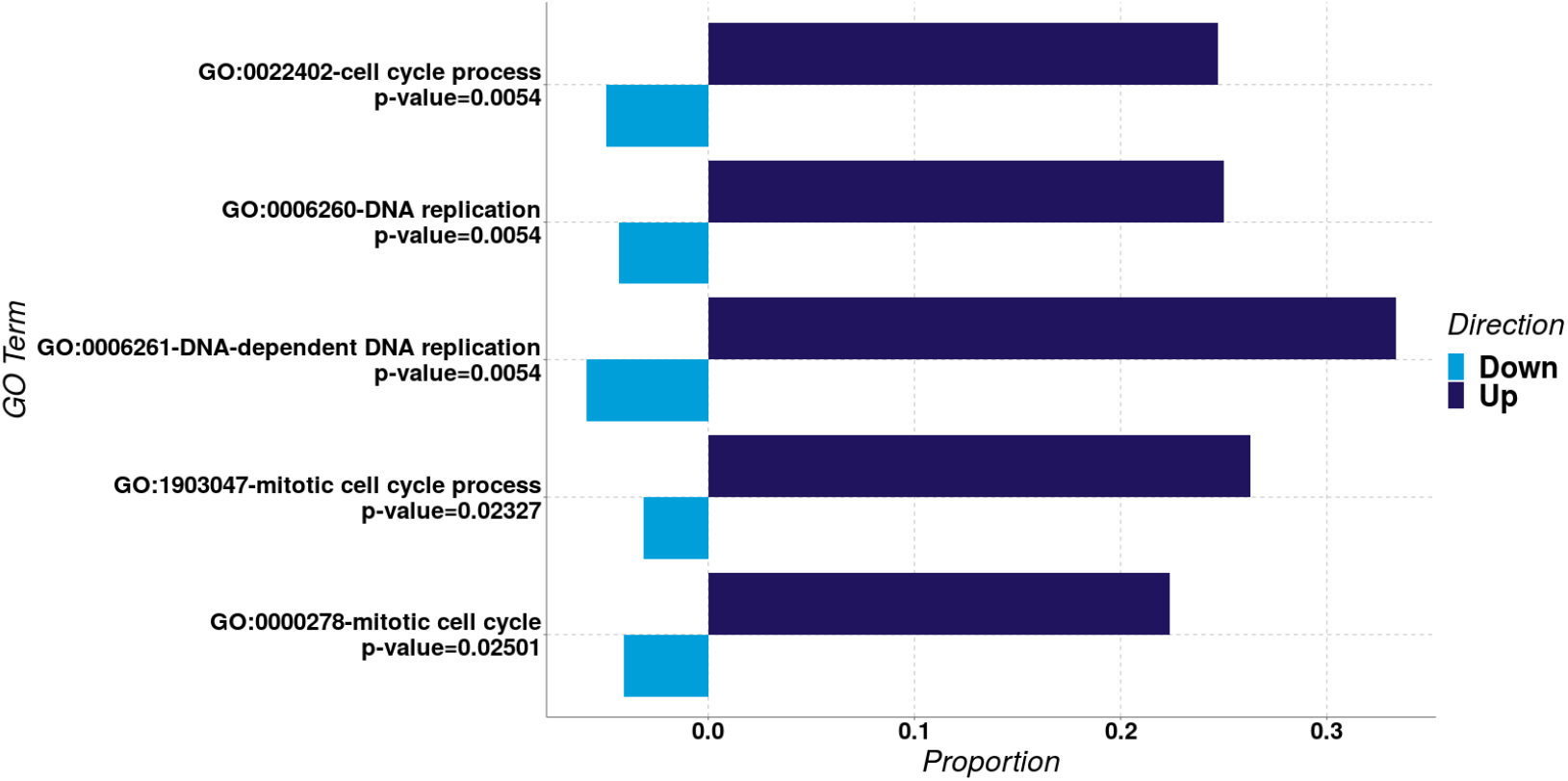
Differential expression (DE) graph showing the enriched pathways identified by clusterProfiler in adult brain. The proportion of individual genes belonging to each pathway is displayed in dark blue for upregulated and light blue for down-regulated expression. The p-value for the enrichment of each pathway is stated below the pathway name.

**Supplementary Figure 2:**
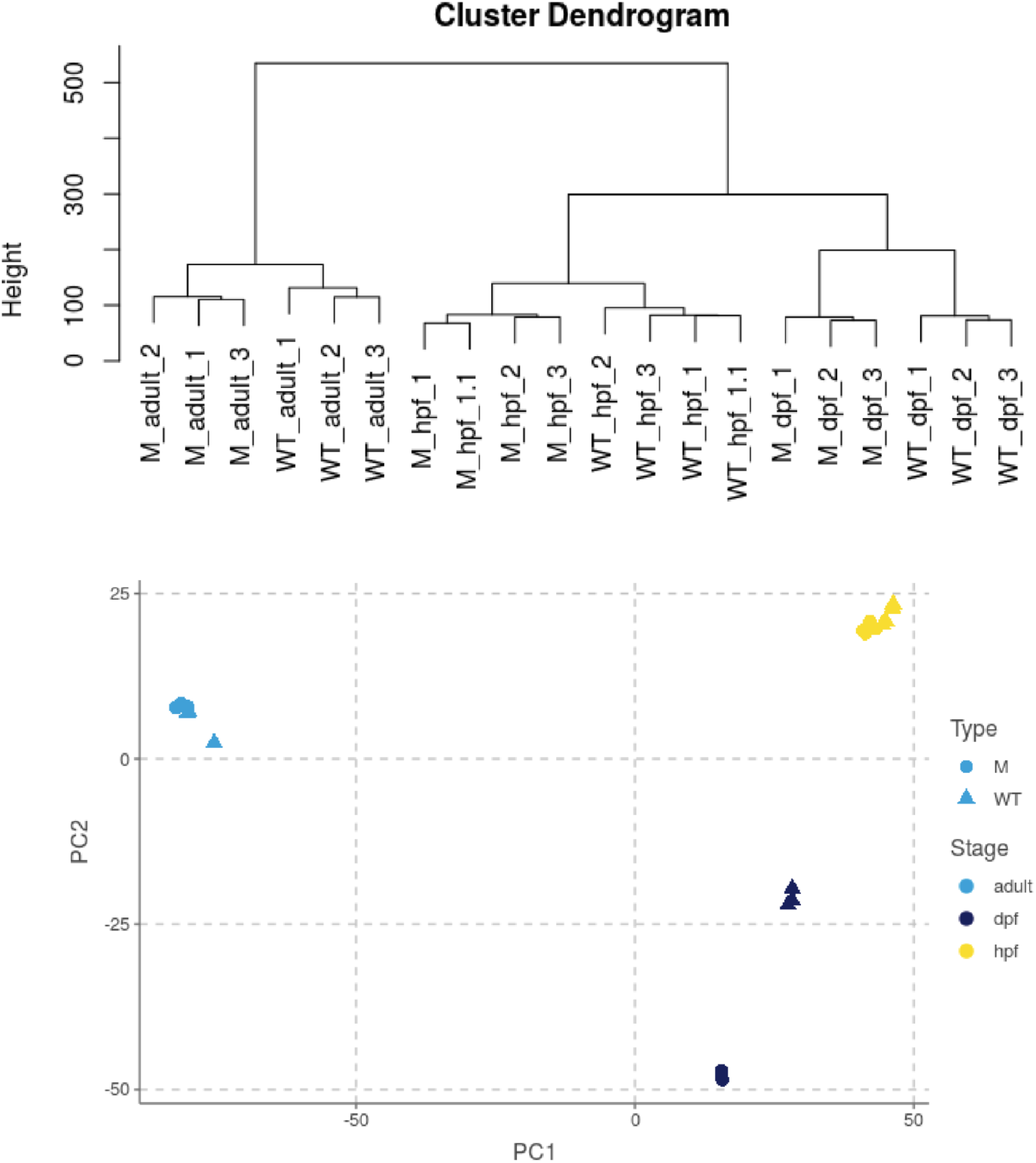
Cluster dendrogram of all timepoints used for microarray analysis showing adult samples clustering separately from embryonic timepoints. M=*rnaset2^ao127^*

**Supplementary Figure 3:**
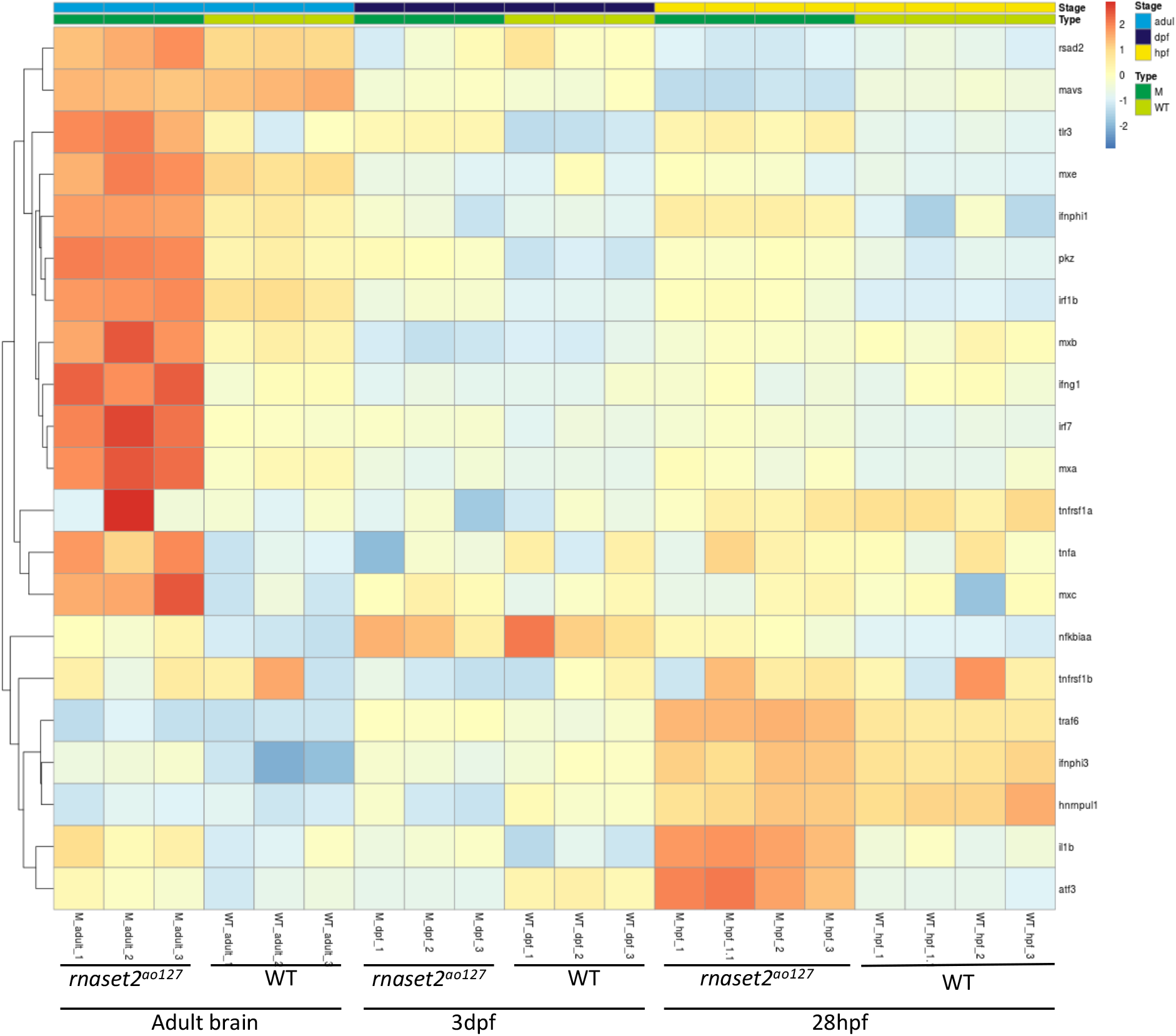
Heatmap of the ‘Response to Virus’ pathway GO:0009615 in all time points

**Supplementary Figure 4:**
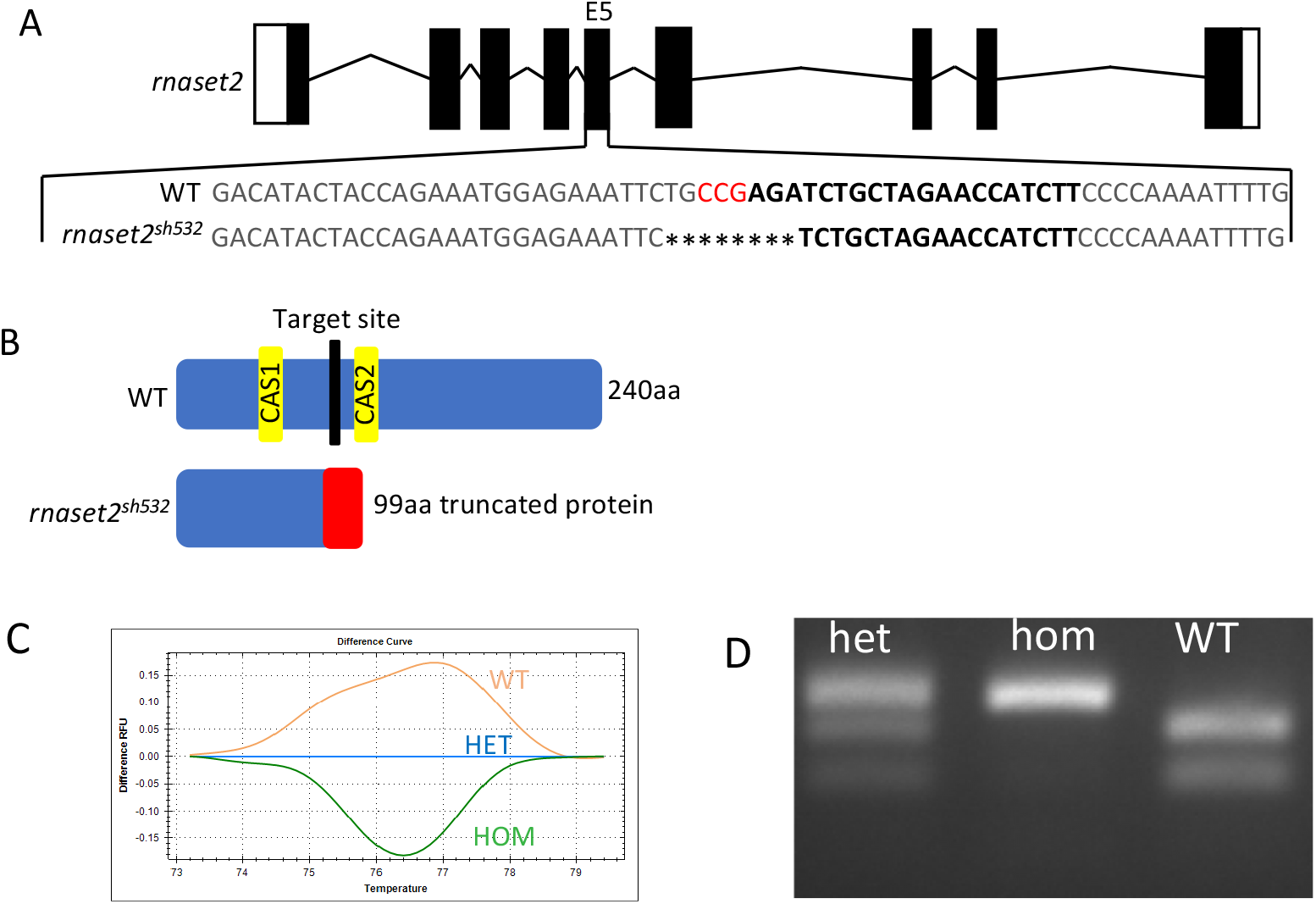
**A.** Diagram of the rnaset2 zebrafish gene with the target gRNA on the WT strand and the selected 8bp deletion in exon 5 (E5) to generate the new rnaset2sh532 allele. **B.** Diagram of the WT rnaset2 protein and the resulting truncated protein in the rnaset2sh532 mutant. **C.** Traces of High Resolution Melt Curve of a WT, heterozygous (HET) *rnaset2^sh532^*, and homozygous (HOM) *rnaset2^sh532^* mutant used for genotyping. **D.** Agarose gel of genotyping of WT, HET and HOM using Mwo1 restriction digest on PCR product flanking the Mwo1 site targeted by the *rnaset2* crRNA on exon 5 of the *rnaset2* gene.

**Supplementary Figure 5:**
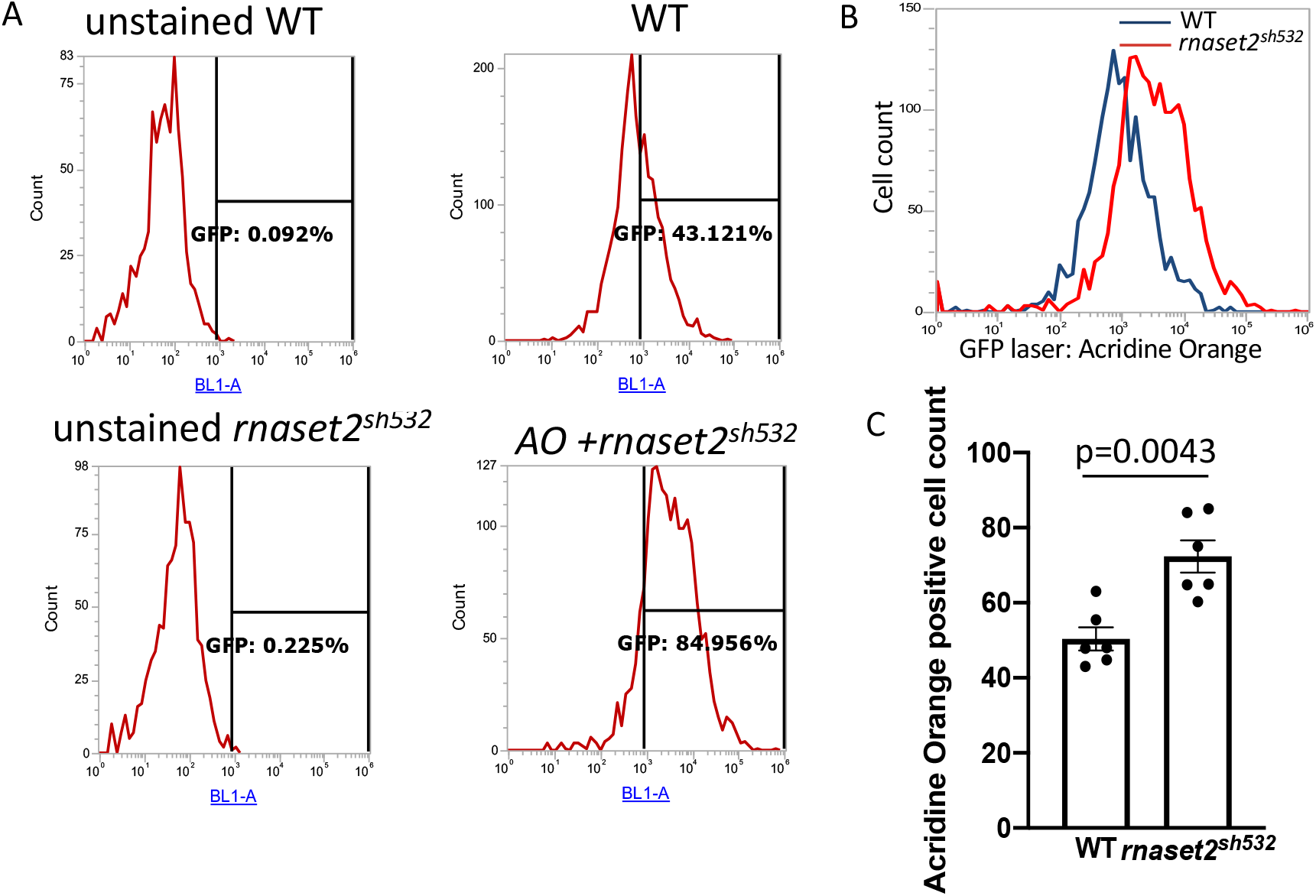
Acridine orange uptake is higher in 5dpf *rnaset2^sh532^* mutant. **A.** Flow cytometry Individual plot of WT and *rnaset2^sh532^* unstained or stained with acridine orange and analyse with gate used to measure acridine orange signal using a GFP laser (BL1A). **B.** Representative overlay of Acridine Orange uptake in 5dpf WT siblings (blue line) and *rnoset2^sh532^* (red line) larvae. **C.** Quantification of acridine orange signal in WT and *rnaset2^sh1532^.* n=6 from 2 biological experiments, two-tailed Mann-Whitney U test p=0.0043.

**Supplementary Figure 6:**
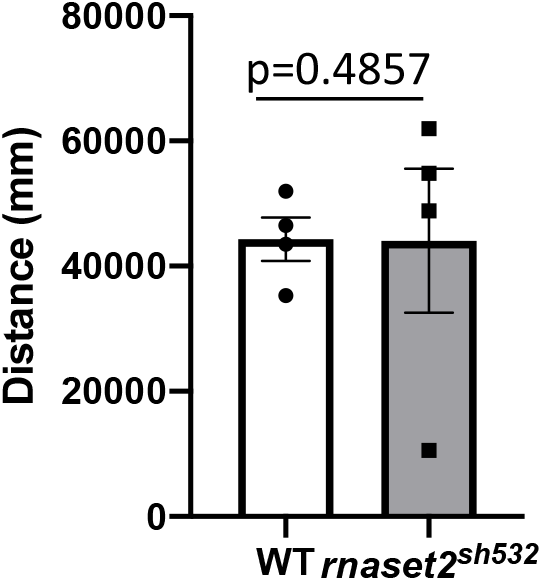
Movement tracking of 8 month old adult fish between WT and mutant for total distance travelled across a 10 minute interval. n=4, two-tailed Mann-Whitney U test.

**Supplementary Figure 7:**
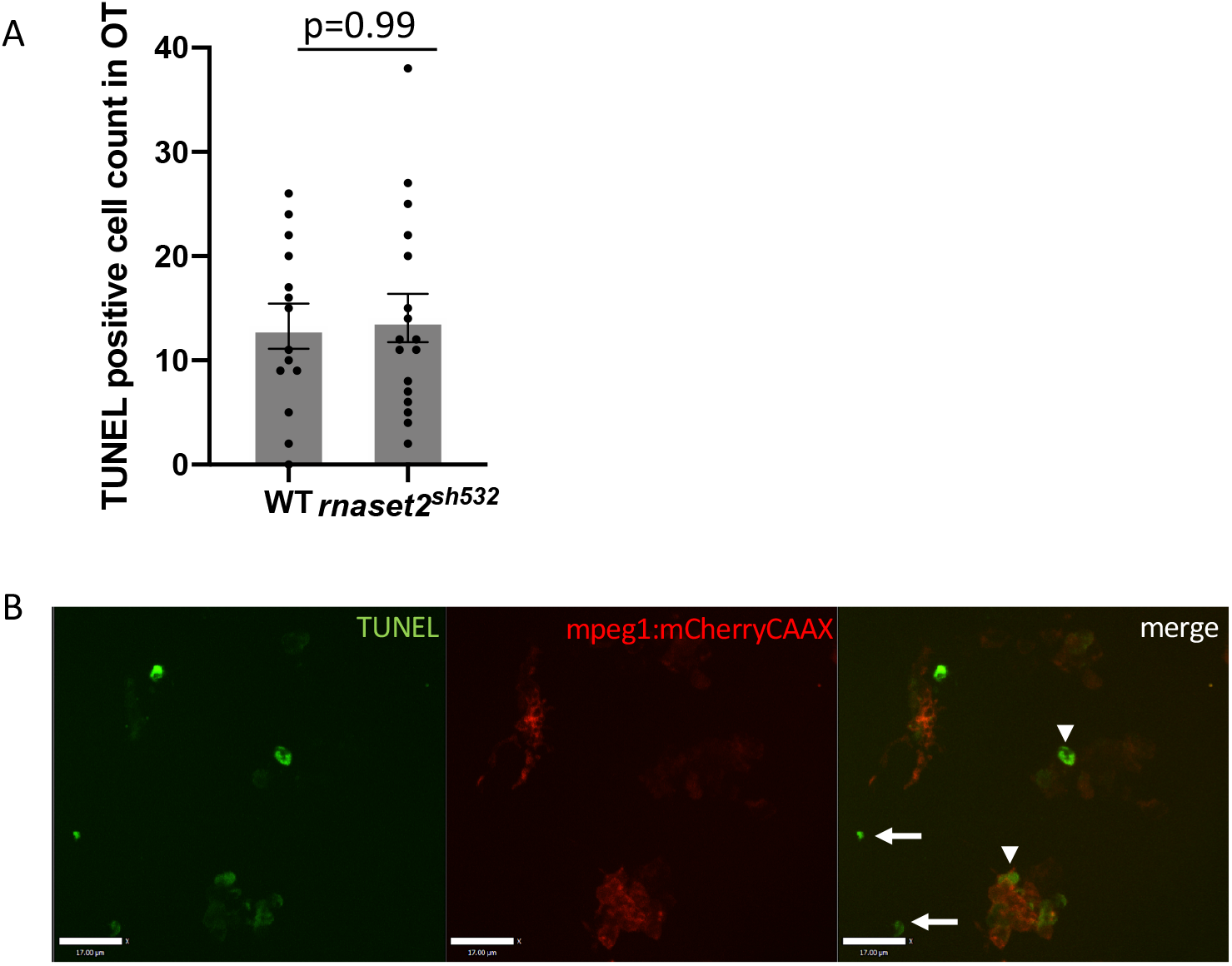
**A.** Quantification of apoptotic cells in the optic tectum (OT) between 3dpf WT siblings and *rnaset2^sh532^* larvae. n=14-17 from 3 independent experiments, two-tailed Mann-Whitney U test. **B.** Representative images from TUNEL positive cells labelled with Fluorescein from 5dpf *rnaset2^sh532^* larvae. Apoptotic cells are either located in the brain tissue (white arrows) or located within microglia labelled in red by the *Tg*(*mpeg1:mCherryCAAX*) line (white arrowheads).

**Supplementary Figure 8:**
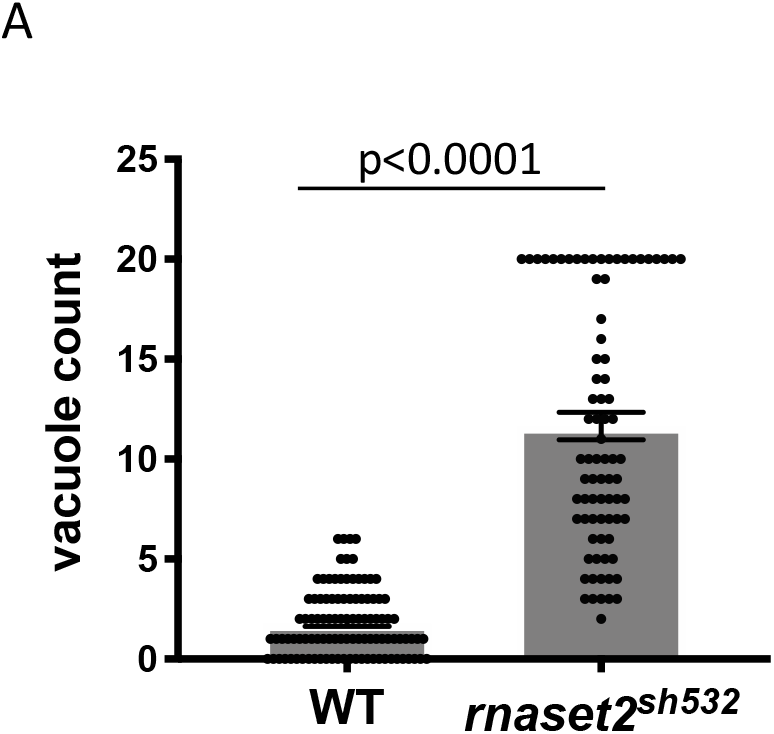
Quantification of numbers of vacuole in individual microglia. Mutant counts were capped at 20 due to the lack of accuracy to identify vacuole when there were too many in each cell. n=80-104 larvae from 3 independent experiments, two-tailed Mann-Whitney U-test p<0.0001

**Supplementary Figure 9:**
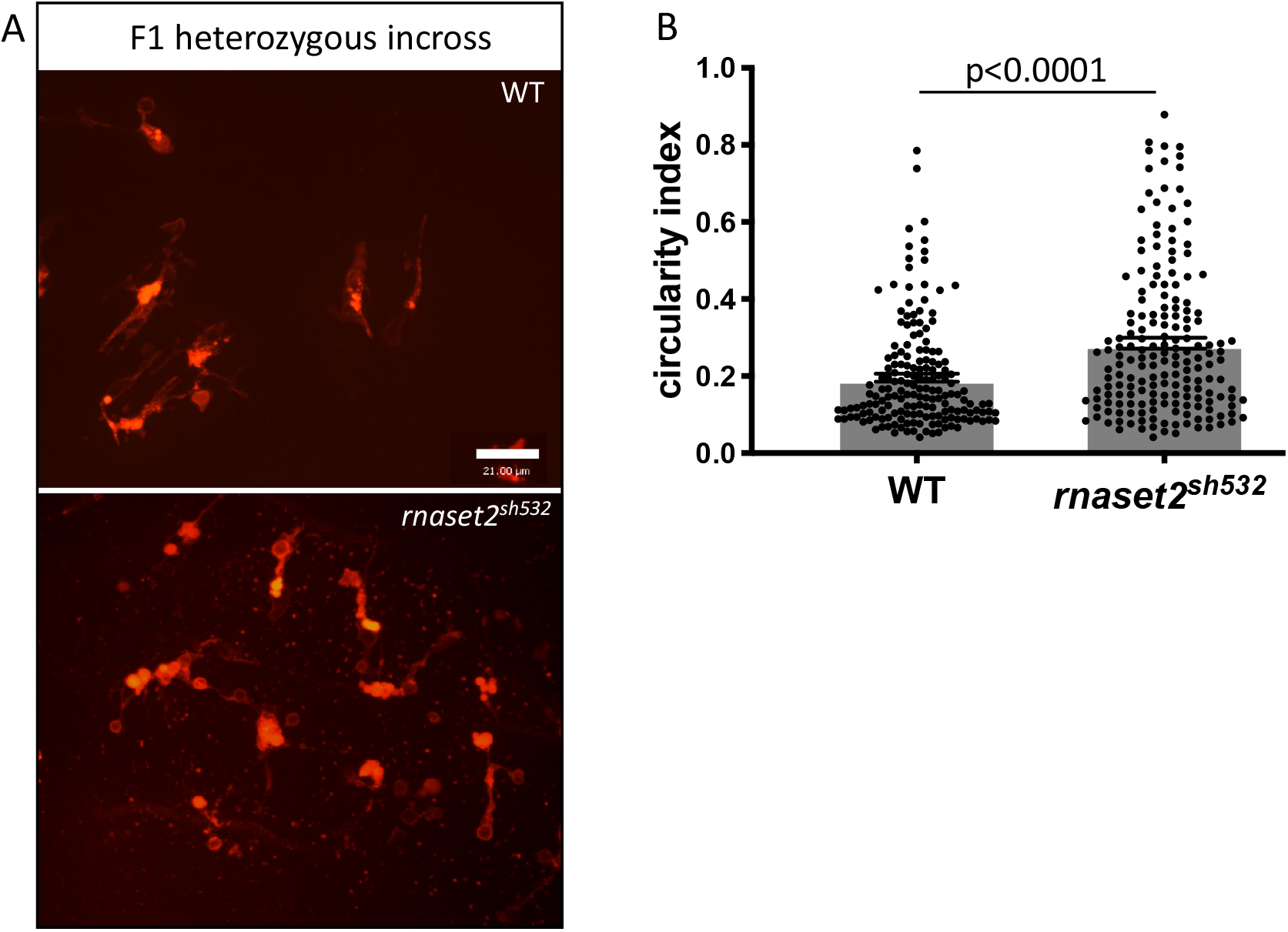
Maternal contribution alleviates defects in microglia morphology. **A.** Representative images of microglia morphology using the 40x objective in 5dpf *rnoset2^sh532^* and WT siblings generated from heterozygous incross. Scale bar 21□m **B.** Quantification of microglia circularity n=173-181 larvae from 2 independent experiments, two-tailed Mann-Whitney U-test p<0.0001

**Supplementary Figure 10:**
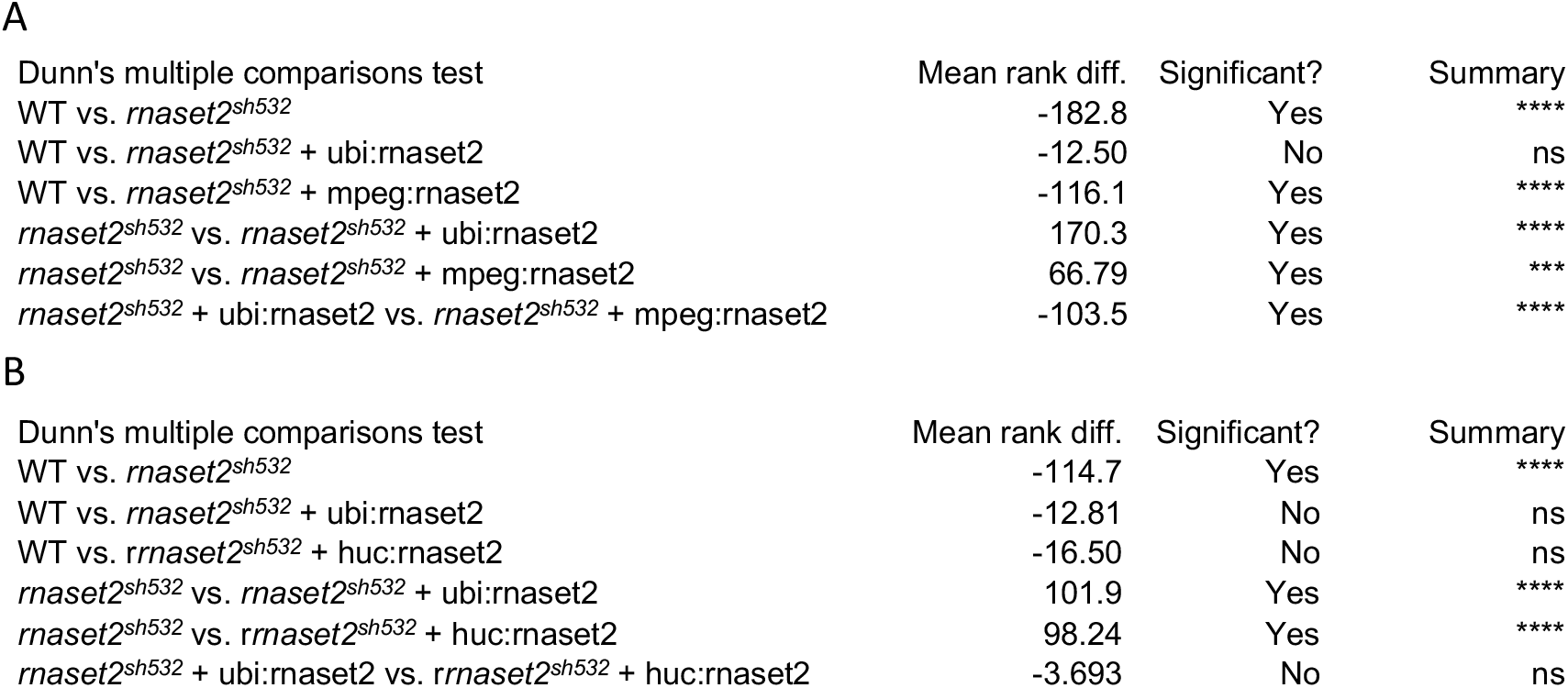
Full results of statistical Krustkal-Wallis test with Dunn’s multiple comparisons for rescue with mpeg:rnaset2 **(A)** and huc:rnaset2 **(B)** from Figure 5.

**Supplementary Table 1.**
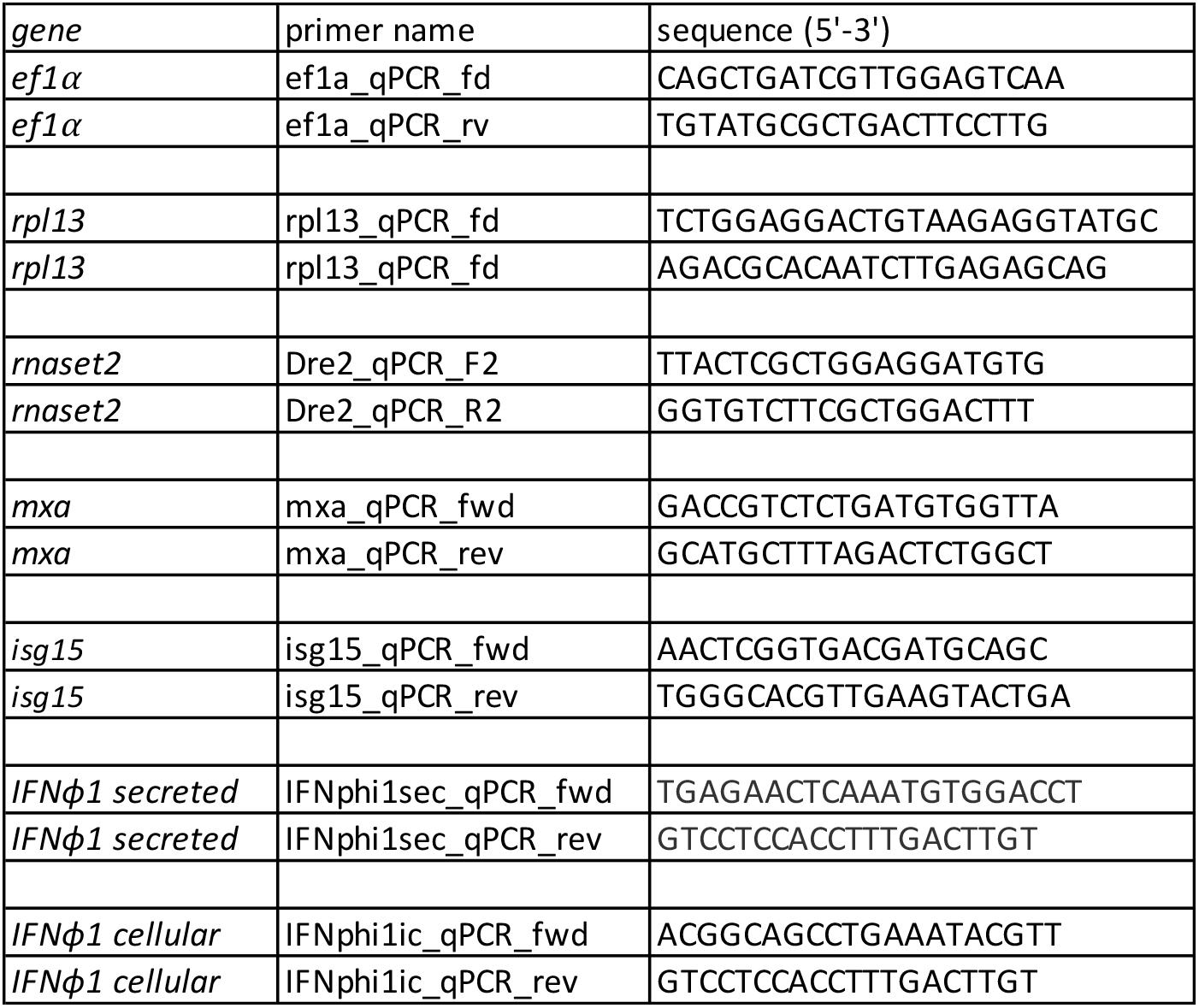

